# Oropouche virus detection in saliva and urine

**DOI:** 10.1101/758839

**Authors:** Valdinete Alves do Nascimento, João Hugo Abdalla Santos, Dana Cristina da Silva Monteiro, Karina Pinheiro Pessoa, Antonio José Leão Cardoso, Victor Costa de Souza, Ligia Fernandes Abdalla, Felipe Gomes Naveca

## Abstract

Oropouche virus (OROV) is an arthropod-borne virus of the *Peribunyaviridae* family, transmitted to humans primarily by *Culicoides paraensis*. It is one of the main arboviruses that infect humans in Brazil, mainly in the Amazon region. We report the OROV detection in the saliva and urine five days after the symptom’s onset. Results were further confirmed by nucleotide sequencing and phylogenetic analysis. To our knowledge, this is the first report that OROV may be detected in the saliva and urine of infected patients. In addition, our results may contribute to the current knowledge regarding the natural history of Oropouche fever.

## Text

The Oropouche virus (OROV) is an arthropod-borne virus, with a triple-segmented negative-stranded RNA linear genome. Each segment is designated according to its size as L (large), M (medium), and S (small). This arbovirus belongs of the *Peribunyaviridae* family, genus *Orthobunyavirus, species Oropouche orthobunyavirus* (https://talk.ictvonline.org/taxonomy/), and two invertebrate vectors have been associated with the urban transmission cycle, *Culicoides paraensis* (Ceratopogonidae), which is considered the primary vector, and *Culex quinquefasciatus* (Culicidae) ^(1)^. Recently, one study reinforced the potential role of *Culex sp*. mosquitoes in the OROV transmission ^(2)^.

The infection with Oropouche virus can result in an acute febrile and exanthematous illness, with symptoms frequently related to other viral infections such as dengue. Oropouche fever cases were reported in several Brazilian states including Amazonas, Acre, Pará, and Mato Grosso, as well as in other South America countries ^(2–9)^.

The Oropouche fever is usually confirmed by detecting OROV genome in the plasma or serum of acutely infected patients, or by specific IgM serology during convalescence ^(1,10,11)^. Nevertheless, recent studies showed arboviruses detection using other body fluids, such as saliva and urine. This was demonstrated to different viral species such as *Chikungunya virus* (CHIKV, family *Togaviridae, Alphavirus* genus) ^(12,13)^; *Dengue virus* (DENV) ^(14)^, *West Nile virus* (WNV) ^(15)^, and *Zika virus* (ZIKV) ^(16–18)^, all members of the *Flaviviridae* family, *Flavivirus* genus. On the other hand, the detection of an Orthobunyavirus in the saliva or urine was not reported to date. Therefore, this study aimed to investigate the presence of OROV in these biological specimens, during the acute phase of the illness.

Between February and June 2016, a total of 352 acute-phase specimens, collected amid 0 to 5 days after symptom’s onset, were sent to Instituto Leônidas e Maria Deane – Fiocruz (ILMD), a research unit of the Brazilian Ministry of Health that was responsible for the laboratory diagnosis of ZIKV during its emergence in the Amazonas State, Brazil. Initially, plasma samples were submitted to RNA extraction using QIAamp Viral RNA Mini Kit (Qiagen), according to the manufacturer’s instructions. Posteriorly, we tested all samples for ZIKV ^(19)^; CHIKV ^(20)^, and DENV ^(21)^, by reverse transcription real-time polymerase chain reaction (RT-qPCR). A multiplex RT-qPCR assay further tested negative samples for Mayaro virus (MAYV) and OROV ^(10)^.

We evaluated the saliva and the urine of five OROV positive patients in plasma, using the same protocol. Subsequently, the OROV positive samples were submitted to conventional RT-PCR targeting L, M, and S segments using a protocol developed by our group. Initially, we performed the reverse transcription reaction using SuperScript IV reverse transcriptase and random primers (Thermo Fisher Scientific). The cDNA was amplified in a reaction using 1.5mM of Mg^2+^, 0.2 mM of dNTPs, 1U of Platinum Taq DNA Polymerase (Thermo Fisher Scientific) and 0.3 μM of specific primers for S and L segments, and 0.5 μM for M segment (Table).

**Table.**
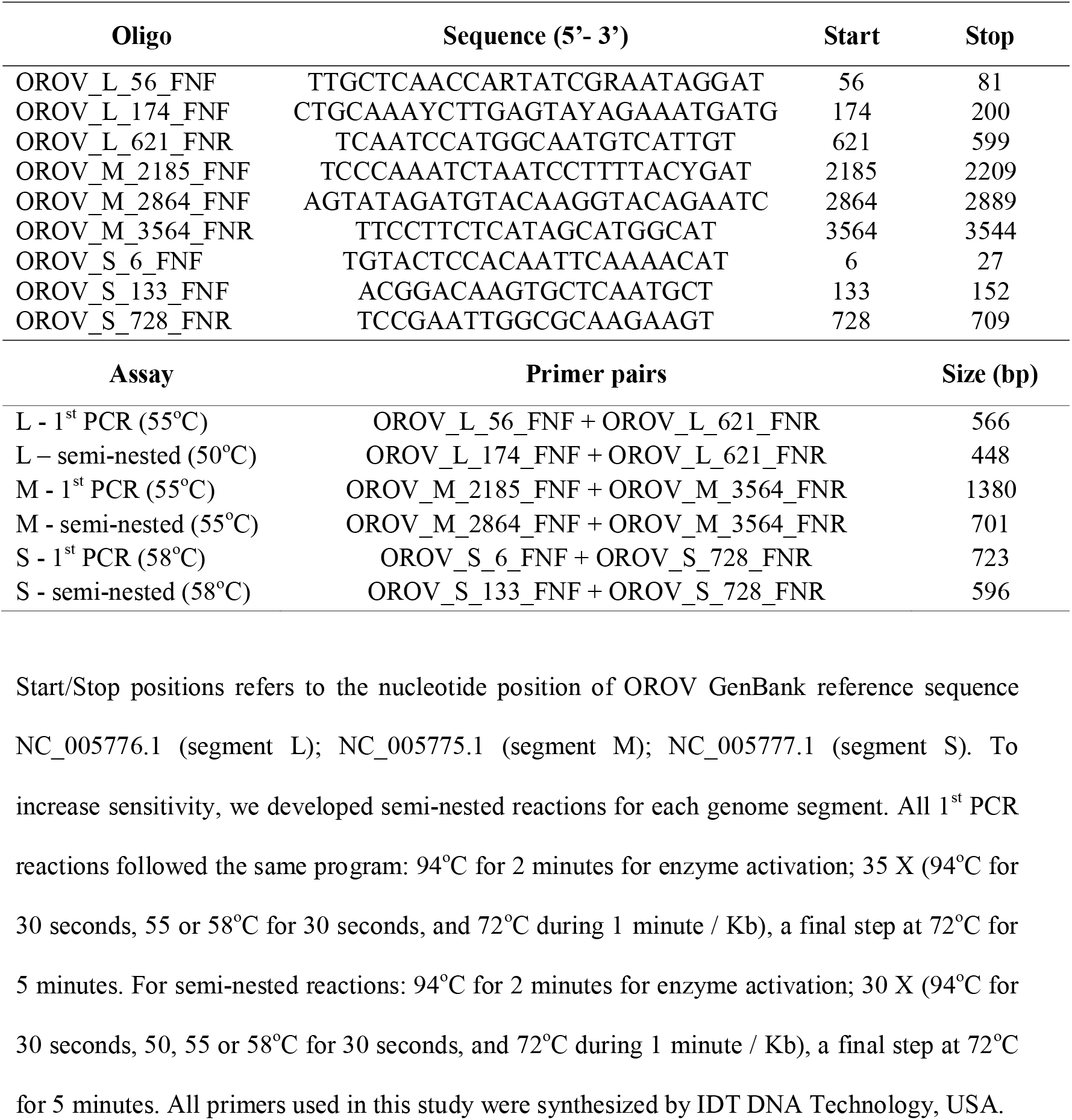
Oligonucleotides designed and used in this study.

The nucleotide sequencing reaction was carried out on an ABI3130 genetic analyzer at the ILMD genomics platform. The data were analyzed using the Geneious software v10.2.6 ^(22)^ for quality check, trimming, and contig assembly. The genome segments sequenced in this study were analyzed together with three different datasets, one for each segment, containing 75 Orthobunyavirus species recognized by the International Committee on Taxonomy of Viruses (ICTV - Virus Metadata Repository: version June 1, 2019; MSL34 - https://talk.ictvonline.org/taxonomy/vmr/m/vmr-file-repository/8287/download), with the full genome records available in GenBank at 01-Jun-2019. All datasets were aligned with MUSCLE (codons) embedded in MEGA X software ^(23)^. Species confirmation was performed using phylogenetic reconstruction by Bayesian Inference (BI) with MrBayes 3.2.6 with two runs and 20 million Markov chain Monte Carlo (MCMC) generations ^(24)^ at CIPRES Science Gateway V. 3.3 (https://www.phylo.org) and maximum-likelihood (ML) with PhyML 3.0 ^(25)^ with Smart Model Selection (SMS) ^(26)^ (http://www.atgc-montpellier.fr/phyml/). All procedures of this study were according to the Ethics Committee of the State University of Amazonas (CAAE: 56,745,116.6.0000.5016).

Among the tested plasma samples, 202 were positive for ZIKV, one for CHIKV, three for DENV, and five to OROV. All OROV positive patients in plasma had saliva and urine further evaluated. One fifty-one years-old female patient (BR_AM_ILMD_0240AOS_2016), living in Manaus, Amazonas State, Brazil, whose samples were collected at 2016-04-11, five days after the symptom’s onset, presented positivity for OROV both in saliva and urine, with Cts values of 31 and 26, respectively. According to the medical records, the patient presents fever, rash, myalgia, pruritus, headache, arthralgia, lymphadenopathy, diarrhea, and vomit during illness.

Partial CDS sequencing was successful for both the L (396bp), M (648bp), and S (555bp) segments and these sequences were used for phylogenetic reconstruction, using a dataset of ICTV recognized Orthobunyavirus species. Both BI and ML phylogeny were evaluated using the nucleotide substitution model GTR+G+I, as selected by the SMS approach. All Bayesian runs reached convergence with an average standard deviation of split frequencies lower than 0.009 and ESS values > 200. For the three genome segments, the sample BR_AM_ILMD_0240AOS_2016 clustered with the OROV RefSeq with high (1.0) posterior probability support (Figure). The same topology was observed in the ML tree (data not shown).

**Figure.**
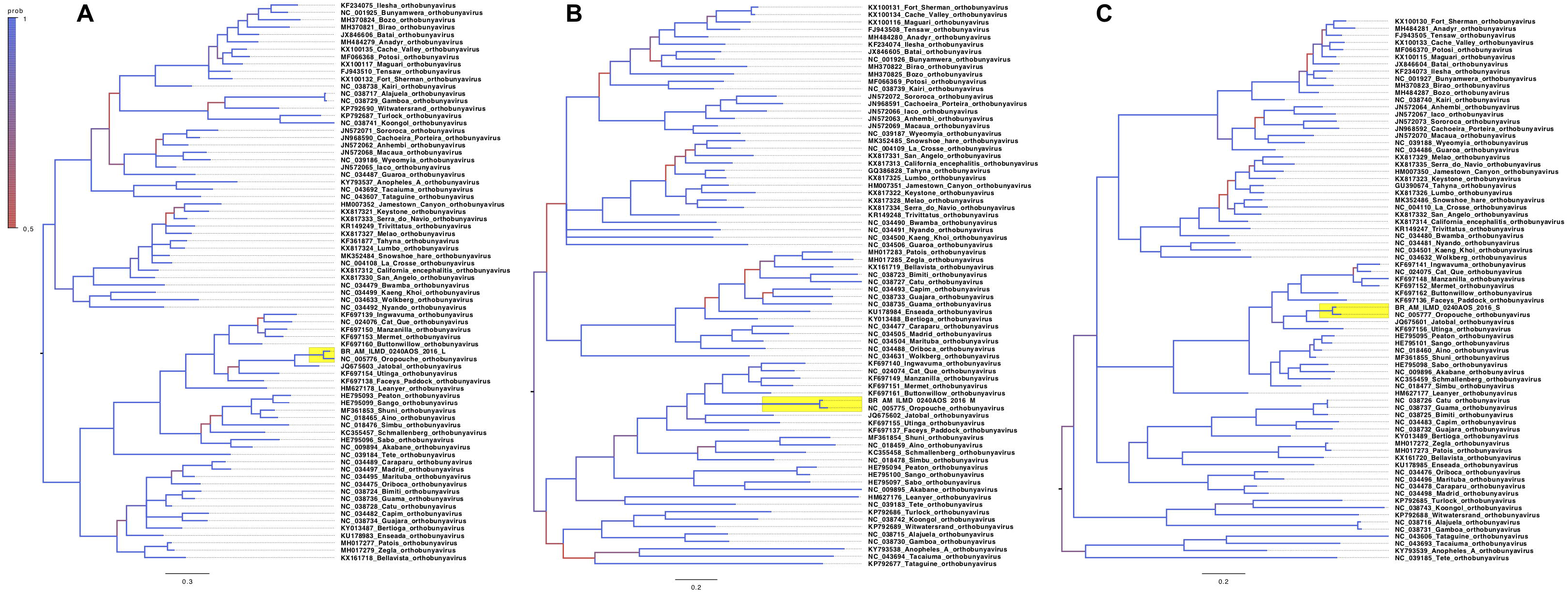
Phylogenetic tree of Orthobunyavirus species. Three mid-rooted Bayesian trees, one for each genome segment, were constructed with MrBayes software v3.2.6 and 76 taxa (the 75 Orthobunyavirus species recognized by the International Committee on Taxonomy of Viruses (ICTV) with complete genome records available in GenBank at 01-Jun-2019 and the sample BR_AM_ILMD_0240AOS_2016 reported in this study). Phylogenetic trees were set mid-rooted, with increased node order in FigTree 1.4.4 for clarity. A color-key represents the posterior probability values of each branch. The clade containing the sequence described in this study is highlighted in yellow, clustered with the Oropouche virus Refseq. The scale bar represents nucleotide substitutions per site. A = L segment tree; B = M segment tree; C = S segment tree.

Several reports have shown that different arboviruses like ZIKV and CHIKV can be detected testing unusual body fluids, such as saliva and urine. Two studies with samples collected from patients infected with ZIKV showed that some individuals were positive only in saliva and not in serum ^(17,27)^. Interesting, our group found similar results during the ZIKV emergence in the Amazonas State, Brazil (unpublished observations). Other reports showed that saliva might be an alternative specimen for CHIKV detection during the acute phase of illness, with positivity raging from 58.3 to 77% ^(12,13)^. However, no previous study reported the detection of a member of the *Orthobunyavirus* genus in these biological fluids.

Therefore, we decided to investigate if OROV, an endemic arbovirus in the Amazon region, could also be identified using the same biological specimens. In the present study, OROV was detected by RT-qPCR in saliva and urine of one patient, whose specimens were collected five days after the symptom’s onset. This result was further confirmed by conventional RT-PCR, followed by nucleotide sequencing and phylogenetic analysis using the ICTV reference database for orthobunyaviruses, which clustered all the three genomic segments sequences obtained in this work with the OROV RefSeqs.

It was beyond the scope of the present study to assess the best human specimen for OROV detection, but it was interesting that we found the higher viral load in urine, when compared with saliva or plasma. Prior studies have also reported the detection of arboviruses in urine. One study with WNV, an arbovirus of the *Flaviviridae* family, reported a higher viral load in urine than in plasma, during the acute phase of illness ^(15)^. On the other hand, two different studies with CHIKV and DENV showed a significantly lower rate of detection when urine was tested during the first days after symptom’s onset, in comparison with samples collected during the second week of illness ^(13,14)^. Altogether, these results suggest that urine may be used as a specimen for different arbovirus detection. Albeit, longitudinal studies, with a more significant number of patients, are necessary to evaluate the potential use of different body fluids for OROV detection.

To our knowledge, this is the first report that OROV may be detected in the saliva and urine of infected patients, showing that these specimens can be employed as alternative sources for the detection of OROV and, perhaps, for other members in the *Peribunyaviridae* family. Moreover, the detection of OROV in urine and saliva strongly suggests that this virus sheds into additional body fluids than blood and Cerebrospinal fluid, as previously reported ^(28)^. Therefore, our results may further contribute to the current knowledge regarding the natural history of Oropouche fever.

### Nucleotide sequence accession number

The partial CDS sequences of OROV isolate BR_AM_ILMD_0240AOS_2016 are available in GenBank, under the accession numbers MN419356 (L); MN419357 (M); MN419358 (S).

## ACKNOWLEDGEMENTS

FGN is funded by Fundação de Amparo à Pesquisa do Estado do Amazonas – FAPEAM call 001/2014 – PROEP / 062.01939/2014); Conselho Nacional de Desenvolvimento Científico e Tecnológico (http://www.cnpq.br, grant 440856/2016-7) and Coordenação de Aperfeiçoamento de Pessoal de Nível Superior (http://www.capes.gov.br, grants 88881.130825/2016-01 and 88887.130823/2016-00) call MCTIC/FNDCT -CNPq / MEC-CAPES/ MS-Decit 14/2016 – Prevenção e Combate ao vírus Zika; Programa Inova Fiocruz, Edital Geração de Conhecimento - VPPCB-007-FIO-18. FGN is a CNPq fellow. The authors thank the Program for Technological Development in Tools for Health-PDTIS FIOCRUZ for use of nucleotide sequencing facilities at ILMD — Fiocruz Amazônia. We also thank the Institut Français de Bioinformatique and France Génomique for the use of the ATGC bioinformatics platform (PhyML), as well as the CIPRES Science Gateway for the use of the Cyberinfrastructure (MrBayes). The funders had no role in study design, data collection and analysis, decision to publish, or preparation of the manuscript.

## AUTHOR’S CONTRIBUTION

VAN, LFA, JHAS, and FGN conceived the study. VAN, LFA, JHAS, and FGN designed the study protocol. VAN, DCSM, KPP, AJLC, VCS, and FGN performed the molecular tests and the analysis and interpretation of these data. LFA and JHAS collected clinical information. VAN and FGN wrote the manuscript. FGN financed the study. VAN, LFA, and FGN critically revised the manuscript for intellectual content. All authors read and approved the final manuscript.

